# Microarray Integrated Spatial Transcriptomics (MIST) for Affordable, Robust, and Comprehensive Digital Pathology

**DOI:** 10.1101/2024.05.31.596759

**Authors:** Juwayria, Priyansh Shrivastava, Kaustar Yadav, Sourabh Das, Shubham Mittal, Sunil Kumar, Deepali Jain, Prabhat Singh Malik, Ishaan Gupta

## Abstract

10X Visium, a popular Spatial transcriptomics (ST) method, faces limited adoption due to its high cost and restricted sample usage per slide. To address these issues, we propose Microarray Integrated Spatial Transcriptomics (MIST), combining conventional tissue microarray (TMA) with Visium, using laser-cutting and 3D printing to enhance slide throughput. Our design facilitates independent replication and customization in individual labs to suit specific experimental needs. We provide a step-by-step guide from designing TMAs to the library preparation step. We demonstrate MIST’s cost-effectiveness and technical benefits over Visium and GeoMx Nanostring. We also introduce ‘AnnotateMap’, a novel computational tool for efficient analysis of multiple ROIs processed through MIST.

## Background

Spatial transcriptomics (ST) has potentially revolutionized the investigation of various disease pathologies, predominantly cancer. It allows study of the positional mapping of cells and their gene expression in a histological tissue section at near single-cell resolution. ST helps delineate the geographical heterogeneity of tumors in terms of cell populations, gene expression signatures, neighboring cell relations, intercellular communication, etc. One of the majorly used spatial transcriptomics techniques is the Visium technology from 10X genomics.

The Visium spatial gene expression slides contain an array of barcoded oligo dT capture primers. Each slide has four capture areas in which the sequencing reaction takes place. Each capture area is surrounded by a fiducial frame and measures 6.5 x 6.5 mm; it contains 5,000 barcoded spots that are 55µm in diameter, providing an average resolution of 1-10 cells. Each spot has millions of primers containing elements required for *in situ* cDNA synthesis, e.g., polyT-tail to capture mRNA. In addition to the PolyT-tail, the capture probes contain a spatial barcode unique for each spot and a unique molecular identifier (UMI) to identify each mRNA molecule. A tissue section, either FFPE or FF of 5 um or 10um thickness, respectively, is placed on the slide, it is first stained by either H&E (hematoxylin and eosin) staining or immunofluorescence staining, followed by synthesis of the spatial gene expression library.

The 10X Visium assay poses 5 major challenges: i) Limited sample usage ii) High cost per sample, iii) Variability among different slide batches, iv) Low level of replication, v) Barcode extraction for data analysis is closely linked with the loupe browser. The loupe browser is a proprietary tool from 10X genomics that is not flexible to accommodate new features to assist with efficient analysis. The visium gene expression slide has four capture areas measuring 6.5X6.5mm sq each. Due to the size limitation of the capture areas, the current protocol recommends using 1 tissue section per capture area. Therefore, only four tissue sections per slide from single or multiple tissue blocks can be used for the assay. 10X Visium is costly; the price of processing a single gene expression slide is approximately 6000 USD, excluding additional sequencing charges. Therefore, many researchers do not readily adopt this technology, especially in low-resource countries. There are two ways in which researchers reduce the cost of imaging-based assays – first, they try to fit in more number of samples in the same imaged area or fiducial frame, and second, they employ computational tools, specifically, machine learning to extrapolate high-resolution data from their low-resolution images. To reduce this prohibitively high cost of spatial transcriptomics so that the technology can be ubiquitously used, we propose to overcome the restriction of limited sample usage and increase throughput per slide through the integration of conventional tissue microarray(TMA) with 10X Visium spatial transcriptomics, introducing a novel approach called MIST (Microarray Integrated Spatial Transcriptomics). We also introduce a novel computational tool named ‘AnnotateMap’ that addresses key challenges posed by the loupe browser (Figure 1C).

**Figure 1:**
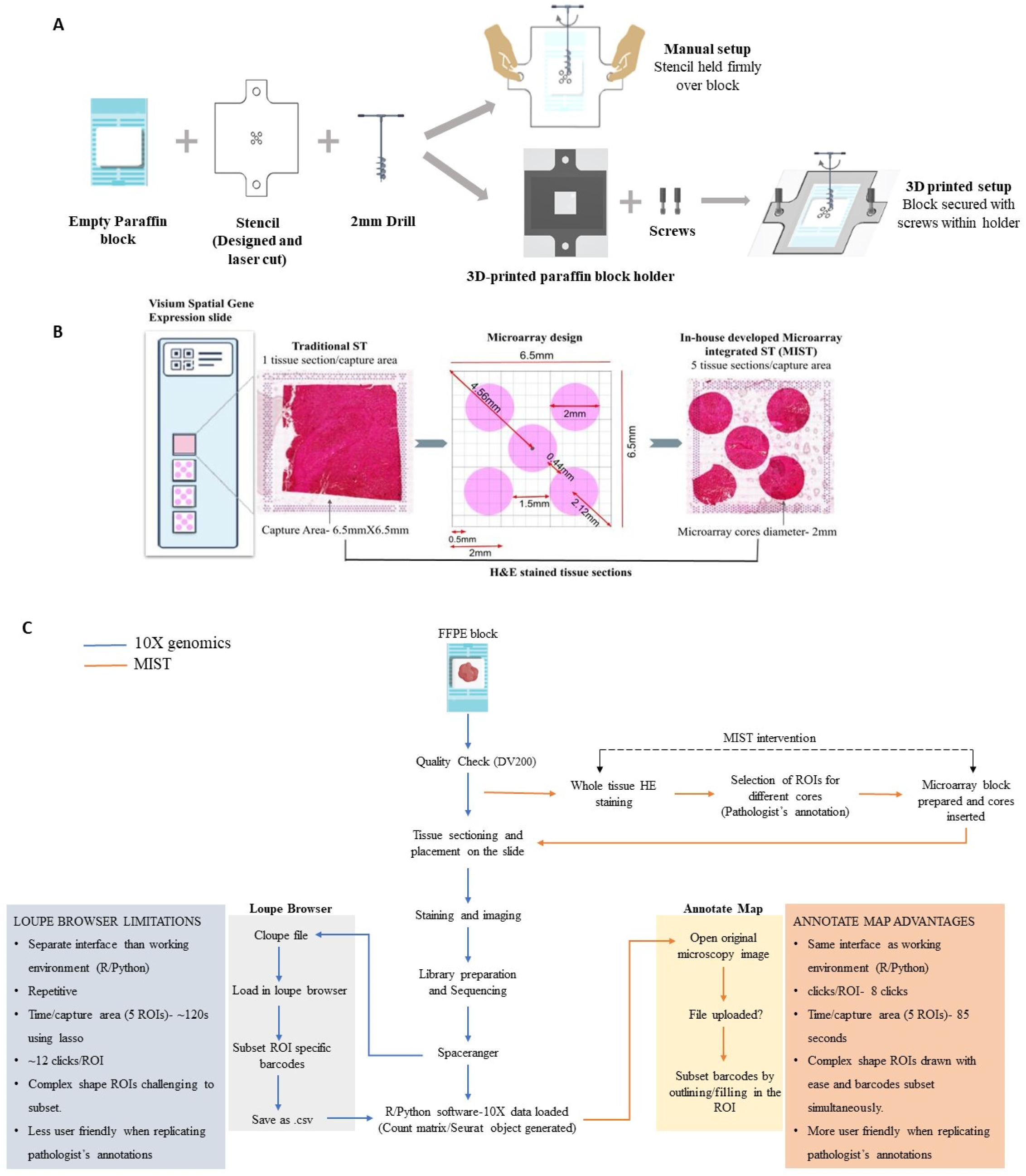
(A), (B) Representative image of traditional spatial transcriptomics and our novel in-house developed microarray approach (MIST). (C) Flow chart depicting the steps of the Visium FFPE assay with MIST, accompanied by a comparative analysis of the 10X Genomics Loupe Browser and our proprietary tool, AnnotateMap.

Tissue microarray was a remarkable invention in pathology in the early 2000s and is still commonly used in hospitals. A tissue microarray uses a single slide to simultaneously assay >1000 minute tissues (1) subsampled from different patient biopsies. It facilitates low-cost multiplexed high throughput analysis of many patient samples and allows the assessment of molecular targets in a cohort under uniform standard conditions, thus removing any variations occurring due to a different slide batch. Tissue microarray enables judicial utilization of irreplaceable tissue resections. This approach leads to multi-fold cost reduction, making the aim of integrating ST into the clinical workflows more feasible.

MIST can facilitate the assay of samples from multiple patients or subsample multiple regions of interest (ROIs) from the same patient where ROIs are distantly located. Multiple sequential sections (replicates) from the same region of interest can also be processed at a reasonable cost. Additionally, it is worth noting that certain regions within the tissue might not offer pathologically interesting insights, such as the presence of a huge blood vessel traversing through a tumor sample, necrosis, etc. As a consequence, these areas might not yield substantial gene expression information. While such occurrences are not uncommon in biological tissues, they may not provide significant information for the specific pathological investigation being conducted. MIST increases the flexibility of the sampling choice, therefore, by deploying MIST, researchers can assay the regions with the most relevant pathological features to interpret variations and capture maximum heterogeneity to ensure comprehensive conclusions.

A substantial amount of tissue is available after surgical resection of entire tumors. However, when dealing with biopsies, the available tissue is often limited, consisting of very small areas. For instance, for metastatic patients, core needle biopsies (CNB) and fine needle aspirations (FNA) are the primary methods utilized in the diagnostic procedure(2), where CNBs are cylindrical biopsies of ≤2mm in diameter (3). MIST can be used to perform spatial assay of multiple (∼20) CNBs in a single slide under identical reaction conditions, which would help prevent potential batch effects and ensure accurate analysis. Similarly, the spatial analysis of multiple FNAs (with a diameter of ≤1mm) can be conducted using the fundamental principles of MIST, as elaborated in the ‘Future Perspective’ section.” MIST can also be employed in pilot studies to generate preliminary data generation for hypothesis building.

While 10X Genomics offers some basic information on TMAs, currently, there is no available guidance on sectioning and tissue placement for Tissue Microarrays. 10X Gemonics recommends using a service provider for preparing TMA blocks and their assessment of TMA compatibility is limited to using 11 mm Visium CytAssist Spatial Gene Expression Slides. Additionally, they don’t offer any guidance on the creation of TMA blocks, troubleshooting, or the placement of TMA tissue sections for CystAssist slides too. Our research article addresses these gaps by offering an in-depth exploration of TMAs. The article includes a comprehensive examination of TMA use, detailing crucial aspects of troubleshooting, tissue type selection, sectioning methods, and the accurate placement of tissues on glass slides. It also discusses various applications and use cases for TMAs, emphasizing their practical utility in research. A notable aspect of our research is the focus on customizing TMAs to suit specific study requirements.

Standard Tissue Microarray (TMA) molds often come with fixed distances between wells, limiting your ability to optimize tissue packing. Although custom TMA designs are an option, they typically require ordering from third-party suppliers. In contrast, our technique is crafted for self-sufficient replication in individual labs. It offers the flexibility to customize and adapt the design according to specific experimental requirements, free from external dependencies. This approach enables labs to efficiently tailor TMAs to their unique experimental needs.

## Results

The proposed protocol consists of four parts: i)3D printing and preparation of the microarray block, ii) Image processing, library preparation, and sequencing, iii)Extraction of core-specific data using the Loupe browser or AnnotateMap, and iv) Data analysis.

10X Visium assay involves tissue permeabilization and probe release onto the gene expression slide, this step may cause diffusion of probes from their original location in the tissue. We 3D printed, optimized, and designed a microarray to accommodate standard circular tissue cores of 2mm diameter from a biopsy, each with enough distance between them to prevent mRNA diffusion across different cores. Additionally, we ensured a minimum distance between the fiducial frame and the tissue cores to prevent overlap with the fiducial frames, as they help in the accurate overlay and alignment of the tissue image with the gene expression data. Keeping these crucial details in mind, we could successfully accommodate five tissue cores in the 6.5 x 6.5 mm capture area. Figures 1A and 1B show the HE-stained tissue with our in-house developed microarray approach (MIST). MIST has only been applied and tested with FFPE tissue samples. The details of the microarray design are mentioned in the Methods section. The flowchart illustrated in Figure 1C outlines the limitations of the Loupe browser that our computational tool, AnnotateMap, is designed to address and overcome.

We have used FFPE tissue sections from resected surgery blocks of a single cancer patient. The whole section H&E staining was done, and 5 different regions of interest (ROIs) to be used for microarray were marked by an expert pathologist; check Methods for details. Figure 2 shows a preliminary analysis of the spatial data generated using the MIST approach.

**Figure 2:**
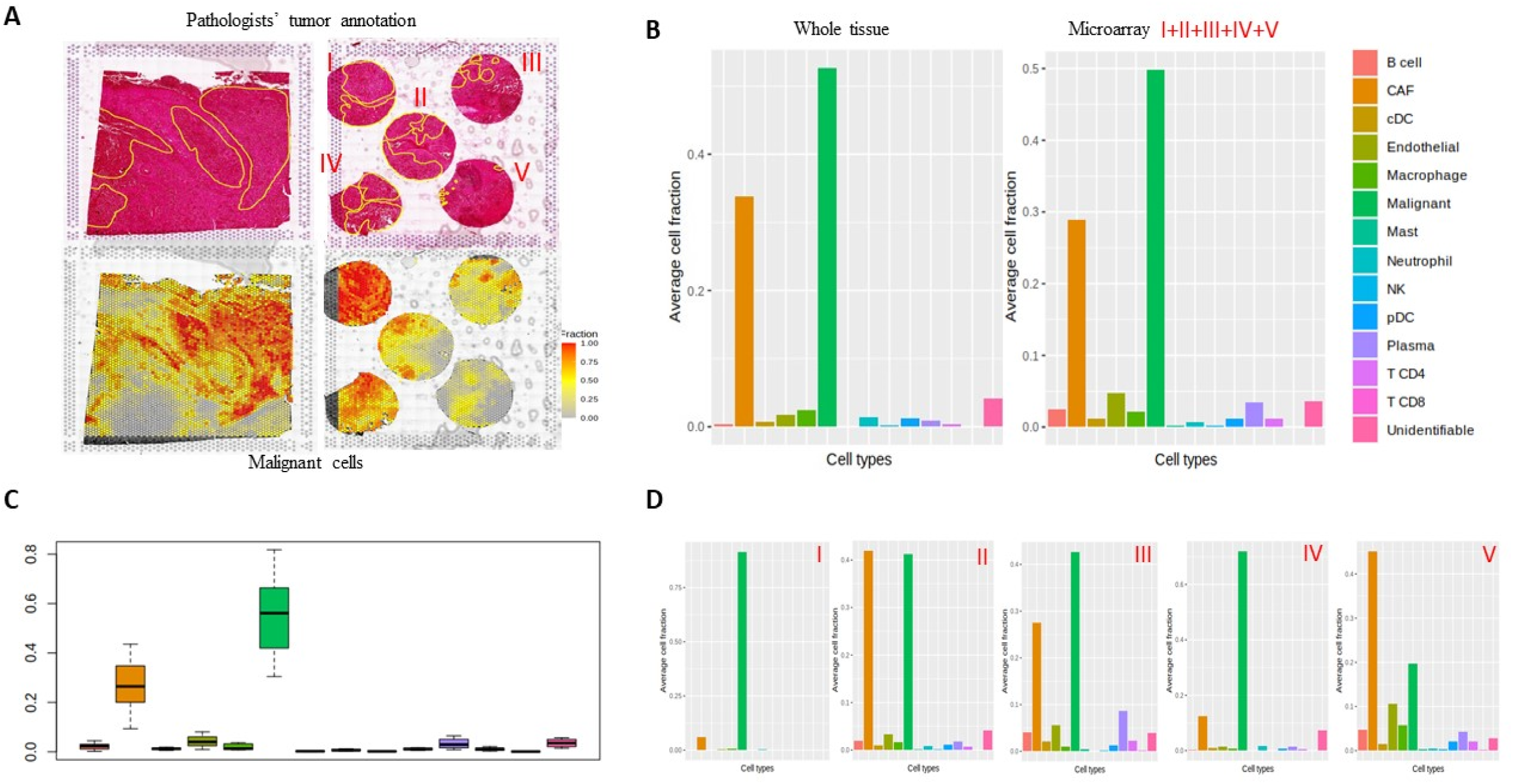
(A) Top left image shows HE stained image of the tissue section spanning the whole fiducial frame. The image in the top right displays a HE-stained image of the microarray derived from the entire tissue. Notably, the five cores in the microarray (labeled as I, II, III, IV, V) were precisely extracted from the identical region from which the whole tissue was initially sectioned. The bottom two images show malignant cell fraction in the two tissue designs. (B) Average cell fractions for different cell types for whole tissue and microarray. For microarray, average fractions from all the five cores are depicted together. (C) Box plot with average cell fraction from any two unique microarray cores combined. (D) Cell fraction from 5 different microarray cores plotted separately.

The major tumor areas annotated by an expert pathologist are shown in yellow contours in the top two HE-stained images of Figure 2A. In the bottom two images, we have overlaid the malignant cell fraction onto the HE-stained image. The malignant cell fraction was annotated using the SpaCET tool, which identifies spots forming malignant cell clusters based on copy number variations (CNVs) and gene expression signatures. We observe that the majority of the areas annotated align with the pathologists’ annotation. The tissue area covered in the microarray design is ∼60% of the whole tissue. Remarkably, we observed that all the cell types recorded in the whole tissue are captured in the microarray design as well. The bar plot in Figure 2B depicts the average cell fraction across all spots for each cell type. Further, we wanted to check if the heterogeneity of a large tissue section spanning the entire capture area can be reflected in a few 2mm cores. To check this, we separately plotted the average cell fraction of the five different cores (Figure 2D) and observed that the cellular heterogeneity can be captured in less than 60% of the entire tissue. Each boxplot in Figure 2C shows the distribution of the specific cell type across 10 different pairwise combinations of the five ROIs. For each combination, the mean of the cell fraction across the two ROIs is plotted. It can be observed that 2/5th of the 60% area of the whole tissue covered in the microarray is sufficient to capture maximum heterogeneity from a whole tissue section. This underscores the need for a judicious selection of Regions of Interest (ROIs) by an expert pathologist, ensuring that the chosen areas accurately reflect the intricate heterogeneity. Customizing the methodology to align with the research objectives is of utmost importance. Depending on the research query, homogenous or heterogenous samples can be selected and assayed. For instance, for studies focused on malignant cells, selecting cancer-specific areas becomes crucial.

The Visium FFPE protocol’s cost is around $6,000 for kits, plus $3,000 for Illumina sequencing (P2 flow cell-400 million reads), totaling about $10,000 per slide or $2,500 per capture area. However, we have made a promising advancement regarding cost optimization. The MIST method significantly reduces the expense of slide processing, bringing down the cost of single sample processing to only 500 USD (1/5th of the original cost, i.e., 2500 USD) or at least 1000 USD if using 2 cores to capture maximum heterogeneity of the tissue.

In order to extract barcode information and efficiently analyze the data from multiple samples, we developed AnnotateMap. Details on AnnotateMap can be found in the methods section.

## Discussion

MIST offers a practical solution to the cost and sample plexity challenges of spatial transcriptomics, making this innovative technique more accessible for comprehensive research in diverse clinical settings. There are tissues with a low cellular density, one such example pertains to the lung tissue. For lung samples, normal lung used as a control contains large alveolar spaces, which do not contribute to significant gene expression information on the gene expression slide as they are essentially empty spaces. To obtain sufficient gene expression data from normal lung samples for comparison with the diseased phenotype, multiple cores from the normal lung must be assayed. Similarly, in the case of high-density cellular regions such as lymph node, a small-sized ROI would capture the distinct cell types. Cancer is a complex molecular disease. Spatial markers for cancer stage identification, drug efficiency, recovery, and relapse once discovered can be utilized to help understand the prognosis of multiple patients together on the same slide with MIST, which would be feasible in clinical applications.

MIST can be employed to gain spatial information from different histology regions as lymph nodes, normal adjacent to the tumor (NAT), tumor, tumor-normal interface, inflammation, vascular inflammation etc. This technology allows for the processing of multiple Regions of Interest (ROIs) from diverse tumor profiles—ranging from cohesive tumor masses to disintegrated tumors or regions with scattered tumor cells—all within a single slide. Such an assay would be very beneficial in use cases where researchers want to explore the cancer evolution in single or multiple patients. While generating spatial data for machine learning or deep learning applications, cost-efficiency is crucial. Here, a larger sample size takes precedence, ensuring an adequate number of samples for effective distribution into training, validation, and test sets. MIST facilitates the simultaneous processing of multiple samples under consistent conditions, effectively mitigating any potential batch effects. Moreover, a larger sample size not only offers a more accurate representation of disease pathology but also captures sufficient variation, mirroring the diversity within a population. Furthermore, MIST stands as a powerful tool to address specific research inquiries. For example, when focusing solely on investigating distinct inflammatory spatial patterns within a particular type of cancer, researchers can meticulously select and analyze multiple inflammatory ROIs, even if these regions are spatially distant. This capacity to selectively target and thoroughly examine specific regions adds a layer of precision and efficiency to cancer research endeavors. In summary, while the initial cost of the Visium FFPE protocol is exorbitant, leveraging MIST allows for substantial cost reduction in slide processing, making this innovative technique an affordable option with single-sample processing at only 2500 USD per sample.

Another spatial transcriptomics technology, GeoMx DSP Nanostring, facilitates the use of tissue microarray(TMA), where multiple circular tissue cores of variable diameter (0.6-6mm) (4) can be fit in the center (35.3×14.1mm)(5) of a glass slide. For DSP nanostring, the region of interest is defined as the area in the tissue from which gene expression data is acquired; an ROI can be further subdivided into different AOIs (segments) with distinct morphologies to obtain separate gene expression profiles(6). Nanostring recommends assaying ≥6 ROIs for a specific region from one patient tissue sample; if ROIs are assayed irrespective of the region type, then ≥24 ROIs are recommended per sample (7). The size range of the ROIs is 10um to 600um (6) in diameter. For RNA analysis, although single-cell resolution (10um) AOI selection is possible, nanoString recommends capturing a minimum of 200 cells from each ROI or segment (7). Any AOI/ROI would give gene expression data as a mini-bulk of cells it encompasses. Hence, as the size of the AOI increases, the resolution decreases, and it would give an expression of all the cells composing the AOI bulk. To compare, MIST based on visium provides a uniform resolution of 50um. As per MAN-10108-01_GeoMx_DSP_Experimental_Design_Guideline.pdf, the usual areas of illumination (AOIs) assayed per ROI are in tens. Such an assay design would not allow a robust understanding of the spatial arrangement inside a tissue; it would only provide information on a limited area (ROI) from the tissue region. It does not come close to the comprehensive analysis MIST provides regarding the complete spatial assay of the core. In our microarray design, the number of spots (analogous to AOIs) analyzed for a 2mm core varies between 400-500.

According to personal communication, the cost of DSP Nanostring is 200 USD for an ROI/AOI of 200um diameter (including sequencing charges). In general, a larger AOI is likely to incur higher costs compared to smaller AOIs, as a larger AOI would require more reagents and higher sequencing depth. Irrespective of the area of the AOI, cost increases as more AOIs are analyzed. If the cost is translated to a 2mm core with 50 AOIs, each having a diameter of 200 micrometers, the estimated cost would be approximately 10,000 USD. In contrast, MIST presents a considerably more economical option, costing 500 USD for analyzing approximately 500 spots (analogous to AOIs), and provides a comprehensive and improved understanding of the spatial dynamics (Table 1).

**Table 1:** Comparison between GeoMx DSP Nanostring and MIST.

| Comparison parameters | GeoMx DSP Nanostring | MIST (VISIUM) |
| --- | --- | --- |
| Setup cost | Requires GeoMx digital spatial profiling instrument (\$295,000) | Benchtop (no investment) |
| Single slide processing cost | 10000 USD for one ROI assaying up to 50 AOIs of 200um diameter | 400-500 spots/ROI- 500 USD. 20 such ROIs can be processed per slide<br>*spots(50um diameter are analogous to AOIs) |
| Choice of sampling | Should be ideally homogenous for a specific ROI | Size of spots is fixed- 50um<br>Heterogenous sampling is ideal- captures maximum heterogeneity |
| ROI analysis e.g. 2mm ROI | Less comprehensive; AOIs, maximum in tens are assayed | More comprehensive, entire 2mm area is assayed |
| Resolution | Larger AOI- more cells-mini bulk of all the cells in the AOI | Fixed resolution corresponding to the spot size(50um) |

In the context of future research, firstly, we propose using MIST for visium slides with larger capture areas and more spots; 11mm x 11mm capture area and ∼14000 spots. Similarly, when paired with BGI STomics, which offers various capture area sizes (50mm², 100mm², 200mm², 36cm², and 174.24 cm²), MIST can potentially enhance the number of samples processed in a single slide. Currently, we have used single-size cores for MIST, we envision an alternative microarray design that uses 2 different sized cores to maximize the use of space within a capture area. In this specific design, we positioned the corner cores at 0.25mm distance from the edge of the fiducial frame. We placed a 1mm core between each pair of corner cores at 0.5mm from each adjoining corner core. This strategic layout allowed us to incorporate a total of nine circular cores comprising two different diameters: five 2mm cores and an additional four 1mm cores. Researchers are encouraged to explore the feasibility of this design for their specific applications. For easy implementation, we have provided the required design files in the supplementary information. Similarly, based on the research question, all 1mm cores, as seen in fine needle aspirations, can also be used to design the laser cut stencil for MIST. This would further allow a more dense packing of the cores and increase the number of samples assayed in a single slide.

## Conclusions

Microarray Integrated Spatial Transcriptomics (MIST) allows flexible selection of diverse regions of interest (ROIs), allowing judicial use of precious tissue biopsies and also reduces the cost of Visium assay by fivefold. We describe a detailed protocol with critical troubleshooting steps to execute TMA integration with Visium ST platform. This work presents the first experimental evidence at the user end with a detailed protocol that performs integration of tissue microarray with an ST platform in a customized manner. MIST democratizes access to spatial transcriptomics for a wider research audience. This approach not only makes spatial transcriptomics more accessible to a broader range of researchers but also allows for the processing of multiple samples under consistent conditions, thereby minimizing batch effects and enabling a comprehensive analysis of disease pathology as discussed in specific use-cases (Discussion). The introduction of AnnotateMap further strengthens MIST’s utility by providing an efficient tool for streamlining the analysis of multiple regions of interest.

## Methods

### Microarray block preparation

We used a 1cm thick acrylic sheet to laser cut a stencil of a larger size (2.5×2.5cm) with microarray cores laser cut as per the proposed microarray design (Figure 1A). Laser cutting technology ensures a precision of 0.1mm, resulting in a highly accurate microarray design. The corner cores are positioned at a distance of 0.5mm from the edge of the fiducial frames, with a separation of 1.5mm between them. Additionally, the center core is situated at a distance of 0.44mm from the corner cores (Figure 1B). A larger stencil, compared to the size of the capture area of the slide, was used to give firm holding space while drilling the microarray block. A blank paraffin microarray block was prepared, and cylindrical cores were drilled out from it with the help of the stencil overlayed on the paraffin block, using a 2mm drill through manual drilling (hand-operated). The stencil can also be printed with a high precision 3D-printer.

### Improved microarray block preparation setup

The described protocol employs manual drilling by positioning the stencil firmly over the paraffin block. However, this manual approach may introduce errors, especially when dealing with smaller than 0.5mm resolutions. To overcome this challenge, we have developed a custom-made apparatus that consists of a 3D-printed mold designed (Figure 1B) to hold a standard FFPE block cassette. Users can place the paraffin block cassette into the mold. On top of this setup, the stencil is positioned, and the entire assembly is secured using screws. Subsequently, the microarray blocks can be created using either a 2mm automated pen drill or manual drilling. The design files necessary for 3D printing the mold and laser cutting the stencil can be found in the supplementary information. We trimmed off the extra wax from the microarray block to achieve a size of about 6.5 x 6.5 mm.

### Tissue selection

This study uses formalin-fixed paraffin-embedded (FFPE) samples from a single cancer patient. RNA extraction was performed with 2-3 FFPE sections of 10um thickness using the Qiagen RNeasy FFPE kit (Catalogue number-73504). The preliminary RNA quality assessment was done on the Bioanalyser to determine RNA fragment size, specifically, the DV200 value (percentage of fragments with ≥200 nucleotides). RNA quality assessment is essential to ensure the success of the downstream library preparation protocol as formalin has a degrative effect on RNA. It is recommended to consult the 10X Genomics user guide and our publication for comprehensive guidance.

### Microarray tissue integration

Simultaneous with the microarray block preparation, tissue sections from whole tissue blocks were HE stained, and regions of interest (ROIs) were annotated by an expert pathologist. After a microarray block with hollow cylindrical cut-outs is prepared, tissue cores from the ROIs are extracted from the whole tissue FFPE block using a standard 2mm cylindrical microarray punch. The cylindrical core is then snug fit into one of the cylindrical cut-outs of the prepared microarray block. Once all five microarray cores were embedded with tissue, the block was incubated at 37°C on a hot plate for 1hr, followed by incubation at 4°C for 15-20min. Post this, the block is trimmed on the microtome for any uneven surface. It is then again put at 4°C for 5min, the blocks are kept at 4°C for long-term storage. We sectioned the microarray blocks 4days after being made, except for one block for which the whole tissue section and microarray section were used on the same slide (samples used in the analysis for this paper). The storage at 4°C aids in allowing the cores to settle effectively within the paraffin block. The optimum temperature and the duration of incubation for microarray block preparation can vary depending on the properties of the paraffin wax. To validate the accurate embedding of ROIs, HE staining was done on sections from the microarray blocks on a plain glass slide and examined by a pathologist.

There are four capture areas (6.5×6.5mm) in the Visium spatial gene expression slide. For the 1st capture area, we performed the spatial assay for a whole section to span the entire capture area of the Visium slide. For the 2nd capture area, once the whole tissue was sectioned and placed onto the slide, an FFPE microarray block was prepared with tissue cores from the same region as the whole tissue. This was done by marking the region on the whole tissue HE-stained slide from which the tissue was sectioned. The marked slide was then overlaid on the tissue to extract 5 ROIs from the same region. The remaining two capture areas (3rd and 4th) were used with the microarray approach from different regions of interest (ROIs), their results are not included in this study.

In the case of whole tissue and microarray sections taken from the same FFPE block (Figure 1A), we adopted a slightly modified microarray block preparation protocol such that the embedded cores are properly settled within the wax block in a shorter time frame. After the TMA block was prepared, it was incubated for 1hr at 37°C, followed by 20min at –20°C. Importantly, the standard visium protocol doesn’t specify if a time delay during subsequent tissue placement on the slide would affect downstream analysis. Due to the study requirements in our protocol, a time gap of 1 hour and 20 minutes was introduced between the first two tissue placements. Through our study, we demonstrate that having such a time gap is safe and does not adversely impact the analysis.

### Tissue sectioning and placement

While tissue sectioning, it is important to ensure an even block surface. If the block protrudes from the upper end in the microtome block holder, there’s a risk of cores dislodging during sectioning. It is important to note down the orientation of the TMA block when placed in the microtome block holder, to further match the orientation of the tissue section floated in the water bath and when placed onto the 10X Genomics gene expression slide.

Sectioning Tissue Microarrays (TMAs) presents unique challenges due to the incorporation of cores from diverse Formalin-Fixed, Paraffin-Embedded (FFPE) blocks. It’s critical to ensure that all tissues undergo uniform processing during fixation and that a consistent type of wax is used for all the FFPE blocks contributing to the microarray cores. This uniformity is key to achieving consistent sectioning quality. To further enhance the sectioning process, it’s advisable to group tissues with similar properties within the same TMA, which helps in maintaining uniform sectioning parameters and improving overall results.

When placing tissue on the 10X gene expression slide, start with the topmost capture area, then proceed to the others. Practice section placement on non-charged plain glass slides with manually drawn capture areas using dummy TMA blocks. Align the top edge of the fiducial frame with any edge of the tissue section floating in the water bath, using a brush for directional assistance, and carefully transfer the section onto the slide. Remember to note the specific edge from which the section is placed on the slide for consistent orientation.

### MIST Assay Troubleshooting: Solutions for Challenges

Microarray block preparation for MIST can present several challenges. Preventing tissue loss while sectioning is crucial, especially while handling blocks with calcified tissue, which can occur due to excess alcohol usage during FFPE block preparation. Calcified tissues become hardened and brittle, making it difficult to section them into thin layers. To mitigate this issue, one must exercise caution and consider either avoiding using such tissue blocks altogether or hydrating them in ice-cold water for 10-15min depending upon the extent of hydration. Hemorrhage commonly occurs during surgical procedures, it may also happen when resected tissue is pressed firmly by forceps prior to fixation (8). Tissue sections with clotting or hemorrhage tend to wash off during staining.

In some instances, the microarray core punched from the whole tissue block may not be uniform or cylindrical due to uneven dimensions of the resected tissue while fixing in paraffin, leading to a loss in the sectioned tissue area. If a significant tissue area is lost, one may punch out the intact cores and reinsert them in another wax microarray block.

If the tissue section exceeds the capture area’s size and risks overlapping with the fiducial frames during the experiment, it can be managed effectively. Place the section on a non-charged, plain glass slide, where 6.5×6.5mm squares are manually marked on the back. This allows for quick trimming of any excess wax. Afterwards, the section can be refloated and then accurately transferred onto the 10X Visium slide. Alternatively, if time permits, extra wax can be trimmed off the wax block.

Additionally, while preparing the microarray block, it is important to drill the paraffin block to the bottom to ensure proper embedding of the tissue cores. Improper drilling will lead to protruding tissue cores, which are at risk of expelling out from the paraffin block during sectioning. This can also happen due to uneven surfaces of the microarray block. In such cases too, another core from a similar pathological annotation may be embedded along with the existing intact cores into a new wax block to avoid loss of space in the Visium gene expression slide. It is advisable to practice microarray block preparation and tissue section placement using dummy blocks before proceeding to the actual experiment slide.

### Library preparation

The FFPE tissue sections placed on the Visium slide were HE stained and imaged using the Leica Aperio Versa 8 microscope at 20X magnification. Library preparation was done in accordance with the 10X genomics protocol for Visium FFPE samples. After library preparation, we used Qubit to measure the library concentration and Tapestation to estimate the size of the fragments. The library concentrations for the four capture areas were 18.2ng/ul, 12.2ng/ul,16.2ng/ul, and 14.7ng/ul. The fragment sizes of the libraries from these areas matched the recommended size by 10X genomics, with peak sizes of 259bp, 270bp, 269bp, and 256bp respectively.

### Data preprocessing

The HE-stained FFPE tissue sections were captured at 20X magnification using the Leica Aperio Versa 8 microscope, which generates images in the.scn format. The size of the.scn files for the entire 6.5×6.5mm capture area is huge, ranging between 1GB and 2.5 GB. In order to make these images compatible with the 10X genomics Spaceranger software, we preprocessed the images using FIJI. Spaceranger requires images in either tiff or jpeg format. To start, we integrated the Bio-Formats plugin into FIJI. The first step is to open the image using the Bio-Formats plugin using the Bio-Formats Importer option. Upon selecting the.scn image, a pop-up window titled ‘Bio-Formats Import Options’ will appear, where only the following four options have to be selected: Hyperstack, Group files with similar names, Use virtual stack, and Default color mode, click OK. Following this, another window named ‘Bio-Formats File Stitching’ will open, keep the default parameters selected. In the subsequent ‘Bio-Formats Series Options’ window, only select the images corresponding to the image of interest, click OK. Once the image opens, the final step is to save the image as either a JPEG or tiff file format.

Post image preprocessing, we performed demultiplexing of Illumina’s base call files (BCLs) into FASTQs by running ‘spaceranger mkfastq’ on Linux with the designated sample indices. Subsequently, ‘spaceranger count’ was run to generate Spaceranger outputs, which were used for further data analysis.

## Data analysis

### Extraction of SpotIDs for each ROI

In order to process the different sample cores from the TMA, extraction of core-specific barcodes is required. Loupe browser from 10X Genomics can be used to extract the spot IDs corresponding to each ROI in the microarray design. cloupe file was loaded on the loupe browser using the ‘Open Loupe File’ option.

Once the.cloupe file is loaded, select the ‘lasso’ option from the top menu, and select an ROI. A pop-up window appears, type the category and cluster name. We have used ‘ST’ as category name for reference. Subsequently, select the spots from all the ROIs using the paintbrush or lasso and save it with a unique cluster name. The recommended approach is to label the Region of Interest (ROI) with the name of the nearest fiducial frame corner, extending from any existing designation, such as triangle, circle, hourglass, or 7-point hexagon. This would prevent any potential confusion that may arise due to image rotation.

The final step is to export the barcodes in a CSV format using the ‘Export Sample-C’ option. The CSV file is then imported to the working environment-R/python for further analysis. The exported barcodes or SpotIDs can be used to subset ROI-specific barcodes from either the Seurat object or the count matrix of a particular capture area. See Figure 3A.

**Figure 3:**
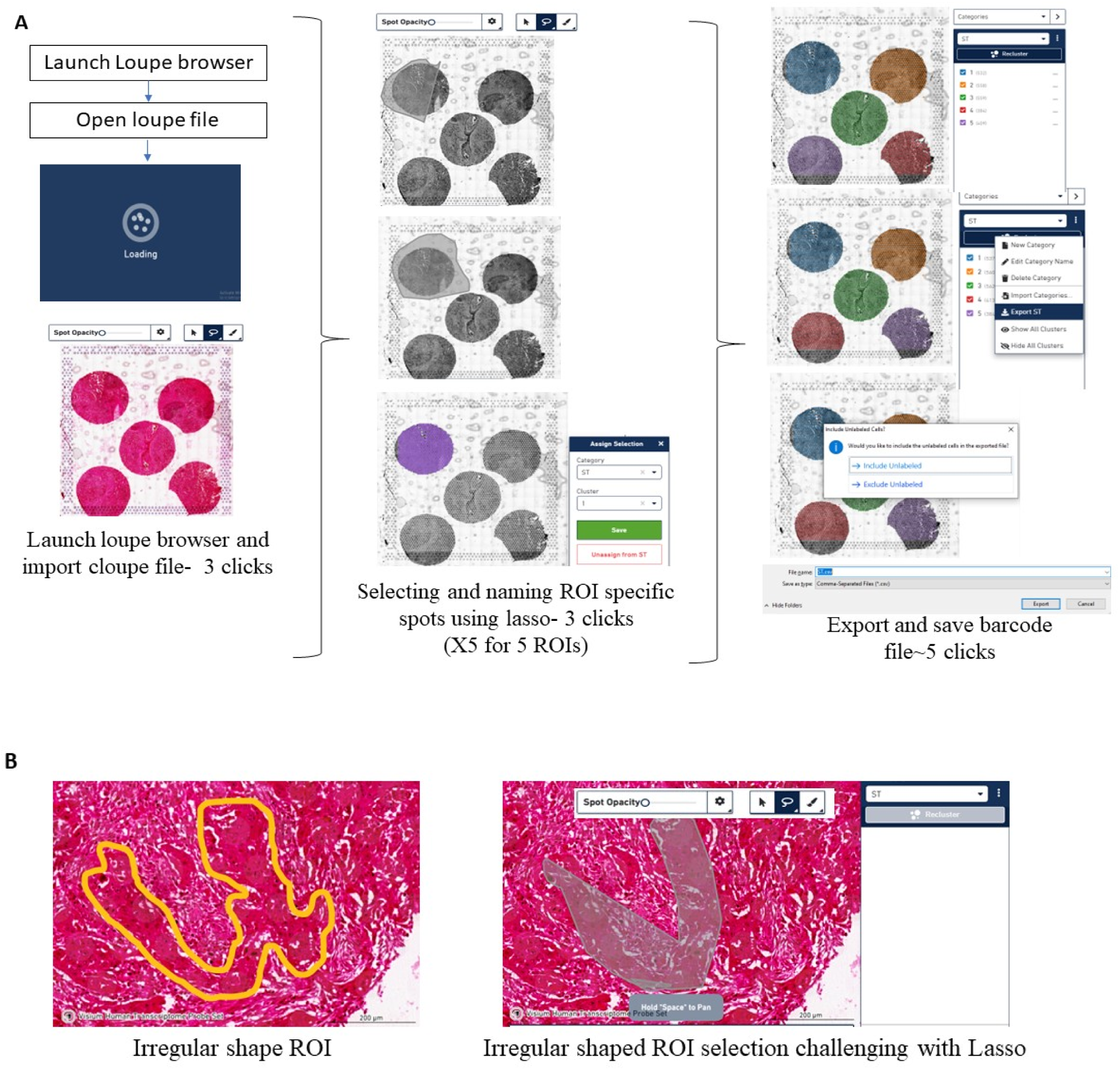
(A) Steps followed in the Loupe browser to subset ROI/core-specific barcodes. (B) Challenge depicted when attempting to select an area with a non-uniform boundary using the lasso or paint tool.

### Overcoming challenges of the Loupe Browser and extraction of core-specific data using AnnotateMap

While the Loupe Browser offers a user-friendly interface, it has certain limitations. The previously outlined steps (Figure 3A) must be performed iteratively to extract barcodes for each Region of Interest (ROI). Time-taken per capture area (5 ROIs) for subsetting barcodes using the ‘lasso’ is ∼2 minutes after sufficient experience with the loupe browser. To put this into context, processing a whole slide would take 8 minutes, and for 10 such slides (common in Spatial Transcriptomics studies), it amounts to 80 minutes. This is further compounded when dealing with huge datasets making the process cumbersome and time-intensive. Given that the Loupe Browser operates as a separate interface, transitioning from R or Python to the Loupe Browser solely for barcode extraction presents additional inefficiencies. Additionally, the need to organize data in two separate interfaces poses a limitation. Having a unified platform for all operations would be highly convenient, particularly when dealing with large spatial datasets. An additional drawback of the Loupe Browser is the limitation in selecting ROIs with non-uniform or complex boundaries using the lasso/paint option (Figure 3B). Manually replicating the pathologists’ annotations again in the Loupe Browser to extract barcodes for further genomic analysis is not very user friendly.

To overcome these challenges, we have developed an efficient and streamlined tool named AnnotateMap. It seamlessly integrates with users’ R/Python analysis workflows, offering a singular interface. Here, barcodes from the 5 ROIs from a capture area can be subsetted by drawing and filling in a boundary around the region of interest, the tool essentially takes 85 seconds to process barcodes from a single capture area with 5 ROIs. Our tool simplifies the process by allowing users to draw boundaries around complex irregular shaped ROIs, making replication of pathologists annotations more user friendly. This also aligns with the intuitive approach typically followed by pathologists.

The integrated Python scripts provide a comprehensive workflow for annotating tissue images and extracting relevant barcode information (Figure 4A). The software.py script utilizes OpenCV to enable interactive annotation of tissue images, offering a resizable display for annotating regions of interest (ROIs) through the drawing of contours. These contours, represented by coordinates, are saved in a CSV file (contours.csv). Subsequently, barcodes.py processes the annotated coordinates from contours.csv and matches them with tissue positional data stored in tissue_positions.csv. It employs a KDTree-based nearest neighbor search to identify the closest tissue elements to each contour point and retrieves their associated barcodes. After removing duplicate barcodes, the script generates a new CSV file (barcodes.csv) containing the unique barcodes corresponding to the annotated tissue regions. This integrated workflow allows efficient annotation and subsequent extraction of specific tissue barcodes.

**Figure 4:**
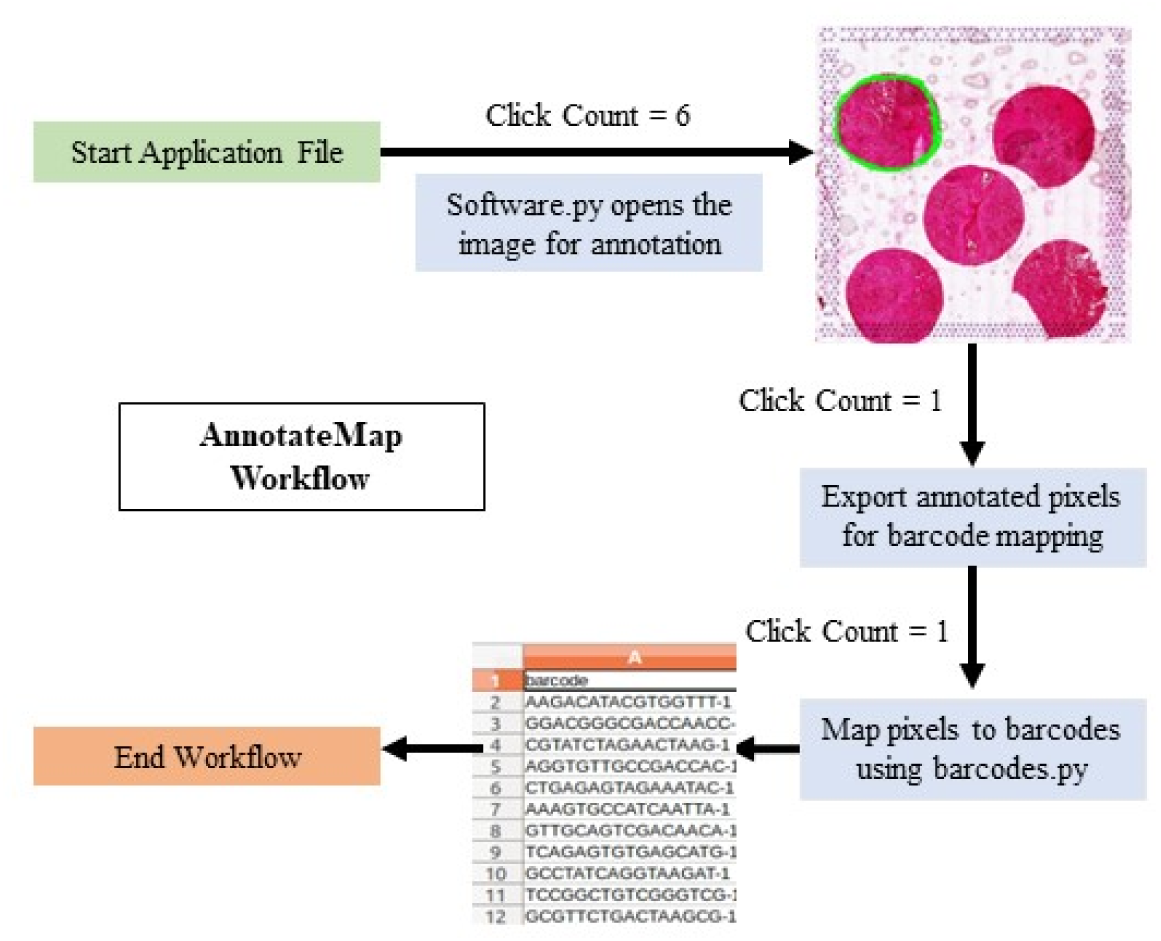
Working of AnnotateMap. More details can be found in our GitHub repository

### Annotation of malignant and non-malignant cell types

Spatial data was normalized using RC normalization on the Seurat object prior to analysis using the SpaCET tool. The Seurat object was converted to the SpaCET object for analysis. 10X Genomics Visium platform offers approximately 1-12 cell resolution in a 50um diameter spot size. In this case, cell type annotation is challenging since multiple cells might contribute to gene expression read out from a single spot. To accommodate this complexity, we used the SpaCET tool (9), which first identifies malignant cells using both copy number alterations and malignant gene expression signatures. Next, it deconvolves the non-malignant cell fraction and adjusts different cell densities in a single spot by employing linear regression, with gene expression profiles of individual cell types identified from various single-cell RNA sequencing (scRNAseq) datasets. The cell fraction in a sample (Figure 2B,2D) was calculated by averaging the specific cell fraction across all spots in that sample.

## Data and code availability

Data related to this study is available upon request. All the supplementary files will be released with the peer-reviewed article. Interested researchers may contact the corresponding author to gain access. Source code for AnnotateMap is available at: https://github.com/Riyu44/MicroarrayProcessor

## Acknowledgment

We would like to acknowledge Abhilash Dasari for his input on the microarray design. Pallavi Gupta and Triveni Shelke for their help in the wet lab experiments. Rajan, for this help in beta testing the working of AnnotateMap. Prakrithi and Nirmal Singh Mahar for their inputs in the analysis discussions.

## Authors’ contributions

Juwayria-designed, optimized, and prepared the microarray apparatus using 3D printing and laser cutting. She contributed to make the microarray blocks, performed the library preparation and sequencing for the Spatial transcriptomics experiment. She performed the data analysis, guided with AnnotateMap, and wrote the manuscript. Priyansh Shrivastava-Lead developer of AnnotateMap and contributed to writing the AnnotateMap section. Kaustar Yadav-Contributed to the preparation of microarray blocks, tissue sectioning, and placement on the gene expression slide. Sourabh Das-Helped with designing the custom microarray design. Shubham Mittal-Contributed to designing the modified version of the microarray setup using 3D printing. Sunil Kumar-Performed the surgery and gave the samples for the study. Deepali Jain-Leading pathologist for the study and contributed to selecting the regions of interest. Prabhat Singh Malik-Involved in ethical clearances, patient sampling and helped in the study design.Ishaan Gupta-Conceptualized the study, guided the data analysis and writing of the manuscript.

## Declarations

### Ethics approval

The necessary ethical clearance was granted by the Institute Ethics Committee, All India Institute of Medical Sciences (AIIMS), New Delhi.

## References

1. Jawhar NMT. Tissue Microarray: A rapidly evolving diagnostic and research tool. Ann Saudi Med. 2009 Mar-Apr;29(2):123–7.

2. Elsheikh TM, Silverman JF. Fine needle aspiration and core needle biopsy of metastatic malignancy of unknown primary site. Mod Pathol. 2019 Jan;32(Suppl 1):58–70.

3. VanderLaan PA. Fine-needle aspiration and core needle biopsy: An update on 2 common minimally invasive tissue sampling modalities. Cancer Cytopathol. 2016 Dec;124(12):862–70.

4. Hernandez S, Lazcano R, Serrano A, Powell S, Kostousov L, Mehta J, et al. Challenges and Opportunities for Immunoprofiling Using a Spatial High-Plex Technology: The NanoString GeoMx® Digital Spatial Profiler. Front Oncol. 2022 Jun 29;12:890410.

5. NanoString [Internet]. NanoString Technologies, Inc.; 2021 [cited 2023 Dec 1]. GeoMx Manuals & Guides. Available from: https://nanostring.com/resources/geomx-manuals-guides/

6. Deshpande N. NanoString. NanoString Technologies, Inc.; 2023 [cited 2023 Oct 16]. GeoMx DSP: What’s in an ROI? Available from: https://nanostring.com/blog/geomx-dsp-whats-in-an-roi/

7. NanoString University [Internet]. [cited 2023 Dec 6]. Spatial profiling of COVID-19 samples: Experimental designs to capture regional heterogeneity with the GeoMx DSP. Available from: https://university.nanostring.com/spatial-profiling-of-covid-19-samples-experimental-designs-to-capture-regional-heterogeneity-with-th/876246

8. Bindhu P, Krishnapillai R, Thomas P, Jayanthi P. Facts in artifacts. J Oral Maxillofac Pathol. 2013 Sep;17(3):397–401.

9. Ru B, Huang J, Zhang Y, Aldape K, Jiang P. Estimation of cell lineages in tumors from spatial transcriptomics data. Nat Commun. 2023 Feb 2;14(1):568.

